# Untangling Sequences: Behavior vs. External Causes

**DOI:** 10.1101/190678

**Authors:** Subutai Ahmad, Jeff Hawkins

## Abstract

There are two fundamental reasons why sensory inputs to the brain change over time. Sensory inputs can change due to external factors or they can change due to our own behavior. Interpreting behavior-generated changes requires knowledge of how the body is moving, whereas interpreting externally-generated changes relies solely on the temporal sequence of input patterns. The sensory signals entering the neocortex change due to a mixture of both behavior and external factors. The neocortex must disentangle them but the mechanisms are unknown. In this paper, we show that a single neural mechanism can learn and recognize both types of sequences. In the model, cells are driven by feedforward sensory input and are modulated by contextual input. If the contextual input includes information derived from efference motor copies, the cells learn sensorimotor sequences. If the contextual input consists of nearby cellular activity, the cells learn temporal sequences. Through simulation we show that a network containing both types of contextual input automatically separates and learns both types of input patterns. We review experimental data that suggests the upper layers of cortical regions contain the anatomical structure required to support this mechanism.

## INTRODUCTION

The sensory inputs to the brain are constantly changing, often several times a second. There are two fundamental reasons why the sensory inputs to the brain change. One is because the body is moving. When the eyes saccade, or the fingers move over an object, or we turn our heads, the patterns of sensory innervations change. The second reason is when sensory inputs change due to external events. For example, listening to music or seeing a bird fly by causes sensory changes that are not caused by our movement. Typically, sensory inputs change due to a mixture of sensorimotor and external causes. For example, as we walk down a street and see a car drive by, the changes on our sensory organs are caused by a mixture of our movements and external causes.

The neocortex receives this rapidly changing input and tries to infer the underlying causes. It must have neural mechanisms to discover which changes in the sensory stream are due to the body’s movements and which changes are due to external causes.

It has been a long-standing hypothesis that information derived from an internal sense of our movements, i.e. efference copy (sometimes referred to as corollary discharge), is involved in this computation (Sperry, 1950; Holst and Mittelstaedt, 1973). The anatomical pathways for efference copy have been studied (Guillery and Sherman, 2002, 2011; Sommer and Wurtz, 2008). Experimental results suggest the cortex forms a predictive model of movements (Duhamel et al., 1992; Nakamura and Colby, 2002; Li and DiCarlo, 2008) and that efference copy is used to form this predictive model (Wolpert et al., 2011; Cavanaugh et al., 2016). However, a detailed neural model of how the cortex uses movement to model and disambiguate sequences is missing.

In two previous papers we proposed models for how pyramidal neurons learn pure temporal sequences (Hawkins and Ahmad, 2016) and how pyramidal neurons learn sensorimotor sequences (Hawkins et al., 2017). We presented these models in isolation. One model learned to recognize temporal sequences by interpreting current input in the context of previous inputs, and the other model learned to recognize physical objects by interpreting current input in the context of a signal derived from the body’s own behavior. The two models had similar components. For example, they both relied on pyramidal neurons with active dendrites and multiple dendritic integration zones. Both models also relied on converting input into a sparse activation of minicolumns, where cells in a minicolumn formed unique representations in different contexts.

Given the sensory input streams arriving at the cortex contain a mixture of temporal and sensorimotor patterns, the cortex cannot rely on two separate mechanisms for modeling the sensory data.

The goal of this paper is to test whether a single neural mechanism can automatically discover what parts of a changing sensory stream are due to movement and which parts are due to external causes, and to learn predictive models of both types of causes simultaneously using simple learning rules. We combined our two previous models into one and through simulation we show that the new single model automatically learns predictive models from both input types. The theory is consistent with a large body of anatomical and physiological evidence. We discuss this support and the model’s relationship to related theories.

## A NEURAL MECHANISM FOR PREDICTION

We have previously described a model of sequence memory (Hawkins and Ahmad, 2016) and a model for sensorimotor inference (Hawkins et al., 2017). Both models rely on the same underlying neural mechanism for learning transitions in the input data and for predicting the next input. The mechanism has two core components: a neuron model that uses active dendrites for prediction, and a model of a cellular layer that learns transitions in the sensory input. In this section we review this mechanism.

**Figure 1A** shows our model of a pyramidal neuron, the HTM neuron. (HTM, or Hierarchical Temporal Memory, is the term used to describe our models of the neocortex (Hawkins et al., 2011).) HTM neurons model the active dendritic properties of pyramidal cells (Spruston, 2008). HTM neurons incorporate multiple dendritic integration zones with different functions. Patterns detected on proximal dendrites represent feedforward driving input, i.e. neurons must detect such a pattern in order to become active. This is analogous to the definition of a classical receptive field in sensory regions.

**Figure 1.**
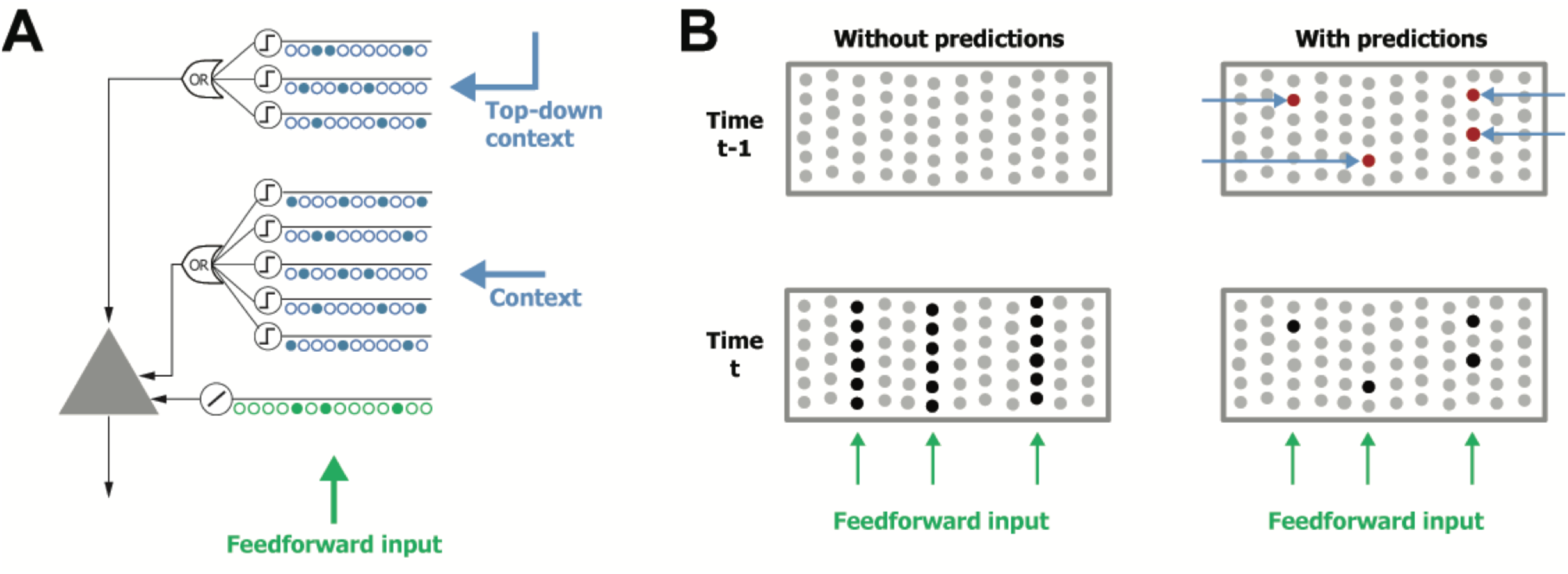
**A.** An HTM model neuron (Hawkins and Ahmad, 2016). Proximal segments (green) detect feedforward patterns and drive cell input. Active dendritic segments (blue) detect contextual input and put cells in a predictive state. **B.** Example showing how a layer of HTM neurons behaves after receiving feedforward input. Top row shows two possible states of the network at time *t* – 1. *Top left*: a layer where no neurons have enough activity on their distal dendrites to cause any predictions. *Top right*: a layer where three neurons detect patterns on their distal dendrites and predict their next input (red). Bottom row shows activity at time *t* for both situations after three minicolumns receive feedforward input. *Bottom left*: with no predictions, every cell in the three minicolumns becomes active. *Bottom right*: the three cells that were in predicted state fire faster and inhibit other cells in their minicolumn resulting in a sparse activity pattern.

Patterns recognized on more distal basal and apical dendrites represent contextual input. These dendrites are composed of multiple dendritic segments, where each segment independently detects a pattern if the number of active synapses on the segment exceeds a threshold (**Figure 1A**). In the HTM neuron model, the detection of such a pattern puts the cell into a “predictive” state that modulates its future behavior but does not drive the cell. A cell in the predicted state anticipates future feedforward input and will fire earlier compared to a cell that is not in this state. The neuron model is similar to (Poirazi and Mel, 2001) except that dendritic segments in our model have a modulatory effect and form a basis for predictions. This behavior models the biophysics of dendritic NMDA spikes, which depolarize cells for 50-100 msecs but do not by themselves cause action potentials (Antic et al., 2010).

A network of HTM neurons arranged into minicolumns can learn to predict future input (**Figure 1B**). Neurons within a minicolumn are constrained to detect the same feedforward input pattern. Inhibitory cells are arranged such that excitatory cells that fire earlier inhibit other cells in the same minicolumn. Thus cells in the predictive state will inhibit the rest of the minicolumn when they become active. Cells that are predicted from both lateral and top-down context are given priority over cells with just lateral predictions. If no cells in a minicolumn are in the predictive state then all the cells become active. This network arrangement closely models the thin vertical minicolumns observed in many cortical regions (Buxhoeveden, 2002). Minicolumns are not strictly necessary; other arrangements work as long as there are small groups of cells with the same classical receptive field and an inhibitory system that favors early firing cells.

The network learns continuously and without supervision. The neuron’s learning rule is based on simple Hebbian-style adaptation: when a cell fires, previously active synapses are strengthened and inactive ones are weakened. There are two key differences with most other neural models. First, learning occurs at the level of the dendritic segment, not the entire neuron (Stuart and Häusser, 2001; Losonczy et al., 2008). Thus cells will form strong connections to contextual patterns on active dendritic segments as long as they are consistently followed by driving feedforward input. Second, the model neuron learns by growing and removing synapses from a pool of potential synapses (Chklovskii et al., 2004). A complete description of the network behavior and learning rules can be found in (Hawkins and Ahmad, 2016).

### Temporal Sequences

In (Hawkins and Ahmad, 2016) we showed that if basal distal dendritic segments form connections to other cells within the same cellular layer, the network becomes a powerful sequence memory. As the feedforward sensory input changes over time, the dendritic segments learn the transitions. If these transition patterns reoccur, the cells predict the possible subsequent inputs. After learning, the activity in the layer becomes sparse as long as the network is making correct predictions. When the network is tracking a sequence perfectly, there is one cell active per minicolumn. Thus sparse assemblies of cells (bottom right panel in **Figure 1B**) become active at each point in a learned sequence. This is consistent with experimental studies of sequence learning in sensory regions (Vinje and Gallant, 2000; Miller et al., 2014). Each cell assembly is unique to a specific point within each sequence, and can be used to predict the next element in the sequence. This is true even for complex non-Markovian sequences, i.e. sequences with long-term temporal dependencies (see Figure 2 in (Hawkins and Ahmad, 2016)).

**Figure 2.**
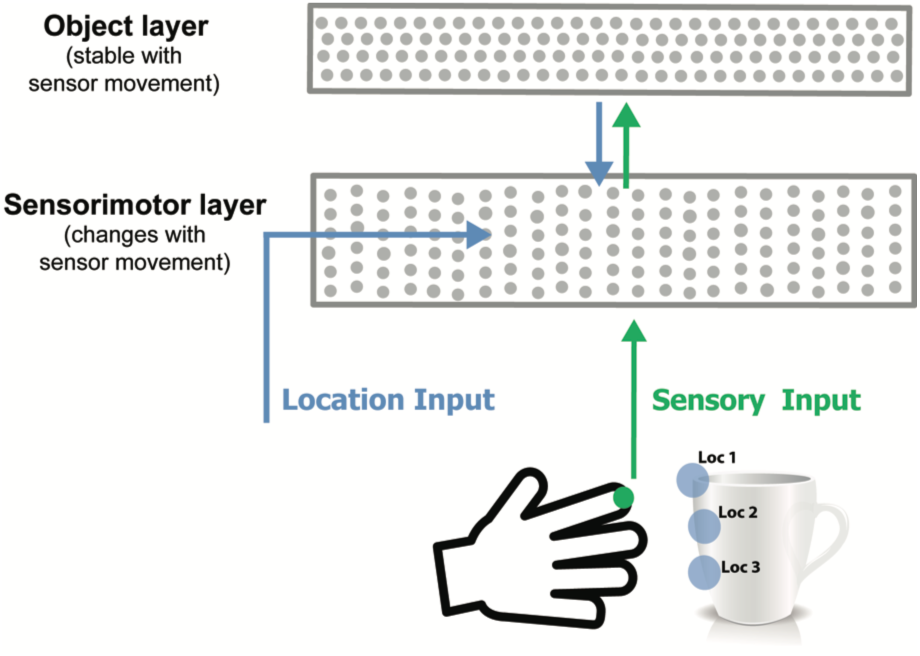
Model sensorimotor network. Adapted from (Hawkins et al., 2017), this figure shows one cortical column only.

In tests with real-world data, a system employing one such layer can make predictions with an accuracy equal to state of the art deep learning algorithms, such as LSTMs (Cui et al., 2016). It addition, due to the continuous learning nature of the model, the system responds to changes in statistics much faster than traditional batch learning systems. The system has also been used for anomaly detection in streaming data applications. In benchmark studies we found that the system performs well compared to other anomaly detection techniques (Ahmad et al., 2017).

### Sensorimotor Sequences

In (Hawkins et al., 2017) we showed that a very similar network can learn predictive models of sensorimotor sequences. As an example, imagine a hand reaching out to touch and manipulate a physical object, such as a coffee cup. The sequence of sensory information is only predictable if you have knowledge of the upcoming movement of the hand, and you know what object is being sensed. Without knowing how the sensor is moving, the sensory sequence will appear random.

To model this, we added a movement generated signal that conveys the upcoming location of the sensor (**Figure 2**). We hypothesized that this signal is computed from an efference copy of the movement command (Guillery and Sherman, 2011). The location vector is fed in as context to the network such that the active dendrite segments on each cell learn to recognize specific locations on the object being sensed. A stable representation of the object is learned in a second cellular layer. This becomes the top-down context to the sensorimotor layer. The activity in both layers becomes sparse as the object is learned and the sensorimotor sequences become predictable. After learning, small assemblies of neurons become active for specific locations on the sensed object. Simulation results show that such a system can form an accurate predictive model of several hundred objects.

In (Hawkins et al., 2017) we also considered the impact of multiple “cortical columns” (as defined in (Mountcastle, 1997) – cortical columns are distinct from minicolumns). We showed that information arriving from lateral columns can greatly speed up inference time. In this paper we limit our focus to just one cortical column, but the results also apply to multiple columns.

## RESULTS

### Combining Temporal and Sensorimotor Sequences

The model we described for sensorimotor prediction is almost identical to the model we described for temporal sequence prediction. The major difference is in the contextual input supplied to the distal dendrites. Given this similarity, we asked if it is possible for a combined system to learn both sensorimotor and temporal sequences, and automatically separate them? A mixed signal is more representative of the data arriving into cortical sensory areas. With externally generated temporal sequences, internal movements are not correlated with changes in the sensory input. Conversely, with sensorimotor sequences, the next input is only predictable if you know the movement that caused the change. We hypothesized that a combined network that had both types of contextual input, and yet had a common learning rule, would be able to automatically learn which contextual information was relevant and correctly predict the next input.

**Figure 3** shows our network model. The lower portion consists of the combined cellular layers. The only difference between the two lower halves is the source of the contextual information. The temporal sequence half receives context from other cells. The sensorimotor half receives context from the motion-derived location input. For clarity, we will refer to the two halves as separate layers, but it could be considered as one layer where some of the cells receive one type of contextual input and the others receive a different type of contextual input. Both the upper and lower layers receive the same feedforward sensory input and share minicolumns. We also include the object layer. It provides top-down context to the sequence layers.

**Figure 3.**
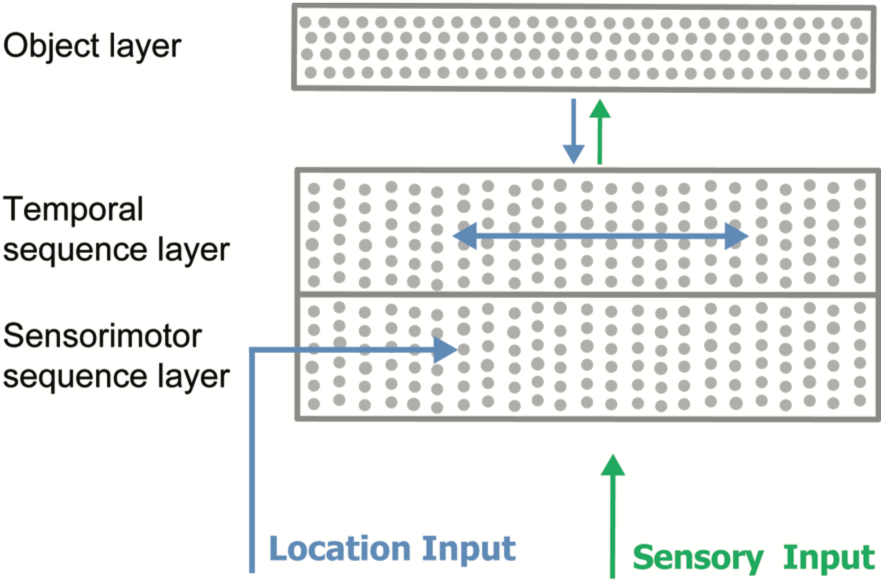
A network with two sequence layers. Both layers receive the same feedforward sensory input vector. The active dendrites of cells in the sensorimotor sequence layer detect patterns in the location input vector. The active dendrites of cells in the temporal sequence layer detect patterns of activity of other cells within the same layer.

In our simulations, both sequence layers had 512 minicolumns with 16 cells per minicolumn for a total of 8,192 cells per layer. The object layer had 4,096 cells. The full network therefore contained 20,480 cells. The feedforward input was represented by a binary vector with 512 components, of which 20 were active at any time. The contextual input for the temporal sequence layer was a binary vector representing the previous activity of the cells within the layer. The contextual input for the sensorimotor sequence layer was a binary location vector with 1024 components of which 20 were active at any time.

The connections in both layers are simultaneously learned and adjusted during the training process. We disabled learning during testing for consistency of results. All network parameters (such as learning rates) are kept constant throughout the simulations and are identical in the two layers (see Methods section below).

### Simulations with Pure Temporal Sequences

We first tested the network on pure temporal sequences. In these experiments, the network was trained on varying numbers of sequences, each of length 10. The elements that comprise each sequence were chosen randomly from a set of possible elements. We presented a randomly chosen location vector for each sequence element.

We controlled task complexity by varying the number of possible sequence elements. The smaller the pool of sequence elements, the more elements are shared between sequences, and the harder it is to make precise predictions. With a large pool of sequence elements (e.g. 1000 features), sequences are more distinct and it is easier to make predictions. One of our goals was to verify that the location input did not adversely affect the ability of the network to predict the next element in temporal sequences. The number of possible location inputs could affect performance. The smaller the pool of location inputs the more likely a specific location could correlate by chance with an element in a temporal sequence. We wanted to test this prediction.

**Figure 4A** shows the behavior of both layers for one example sequence. In this experiment, the network was trained on 5 sequences. The feature and location input pools were 10 and 100 vectors, respectively. This led to sequences with many shared elements (the mean number of input elements shared between two sequences was 9). The figure shows that the sequence layer is able to disambiguate and predict the input sequence. After the first two inputs, it predicts the correct set of 20 minicolumns for each subsequent step in the sequence.

**Figure 4.**
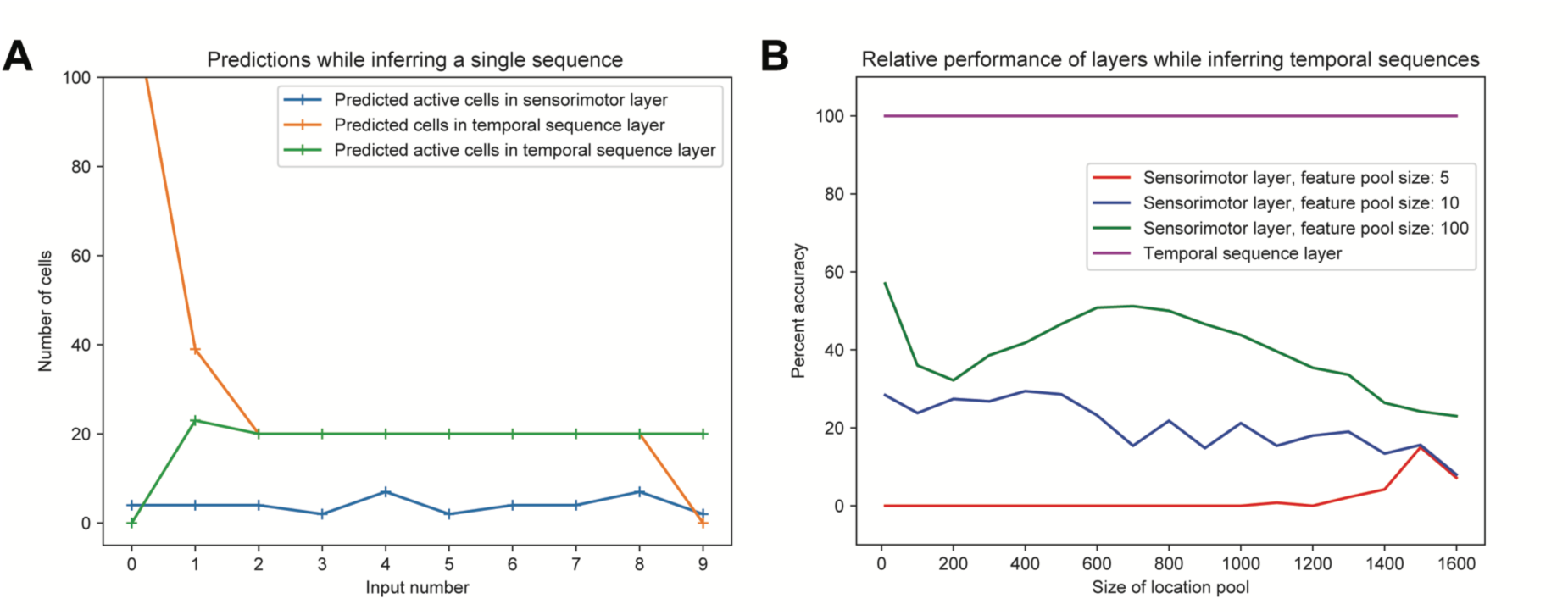
**A.** Network behavior while inferring a single sequence. "Predicted" cells refer to cells with a contextual pattern detected on at least one distal segment. "Predicted active" cells refer to predicted cells that subsequently detected a feed forward pattern, i.e. they correctly predicted future activity. Since 20 minicolumns are activated at each time step, a prediction of 20 cells which are all subsequently activated would constitute perfect prediction. The temporal sequence layer generates more than 20 predictions during the first two inputs due to shared prefixes. The predictions rapidly narrow down and become perfect. As expected, the sensorimotor layer fails to make a significant number of correct predictions. **B.** Relative accuracy of sensorimotor layer as a function of number of locations and feature diversity. Each point is the average accuracy across all trained sequences. For reference, we have included the performance of the sequence layer, which achieves perfect accuracy in all conditions.

This is consistent with our previous results (Cui et al., 2016). The sensorimotor layer on the other hand had a very small number of correct predictions, something we consistently observed with many different parameter combinations. Note that both layers receive identical feedforward input patterns.

To gain a better sense of the behavior of the sensorimotor half of the network during temporal sequence inference, we systematically varied the size of the feature and location pools. To quantify performance, we calculated how often the set of representations in the sensorimotor sequence layer was unique for a temporal sequence. This is identical to the measure used in (Hawkins et al., 2017), where the object layer converged on a stable and unique representation if the sequence of representations was unique. We fixed the number of sequences learned at 50, therefore chance performance was 2%. **Figure 4B** shows a plot of accuracy while varying the size of the input feature pool and the location pool. With a small feature pool, the sequences are highly ambiguous and accuracy is essentially at chance, regardless of the number of locations. As the feature pool increases, the sequences become more distinct and the sensorimotor layer performance increases somewhat. However, its performance is never close to the performance of the temporal sequence layer, which is always at 100%. The underlying reason is that the sensorimotor layer is never able to consistently associate locations with sequence elements and create unique representations.

### Simulations with Sensorimotor Sequences

Next, we tested the network with sensorimotor sequences. As in (Hawkins et al., 2017), we trained the network on a sequence of features and location inputs generated by randomly exploring objects. Each object consists of 10 features, where each feature is assigned a corresponding location on the object. Since features on the objects were explored in random order, there is no inherent consistency in the sequence of inputs. The next input is only predictable if the location is known.

We controlled task complexity by varying the size of the feature pool and the training set size (number of objects). With a small feature pool, the diversity of objects is restricted, and the transitions between locations are more likely to repeat. Thus, the exploration of simple objects will start to look like temporal sequences. We kept the number of locations fixed at 100 since this does not affect the temporal sequence layer. We have previously verified that the sensorimotor layer works well with a wide range of location pools. As in the previous simulations, both layers receive identical feedforward input patterns.

**Figure 5A** shows the behavior of both layers for one example object. In this experiment, the network was trained on 50 objects, with a feature pool of 100 vectors. The figure shows that the sensorimotor layer is able to correctly predict each input feature, consistent with our previous results (Hawkins et al., 2017). This verified that the temporal sequence layer did not interfere with sensorimotor inference. The temporal sequence layer however generated a very small number of correct predictions (predicted active cells). The layer does occasionally make a reasonable number of predictions, but these predictions are mostly random and incorrect.

**Figure 5.**
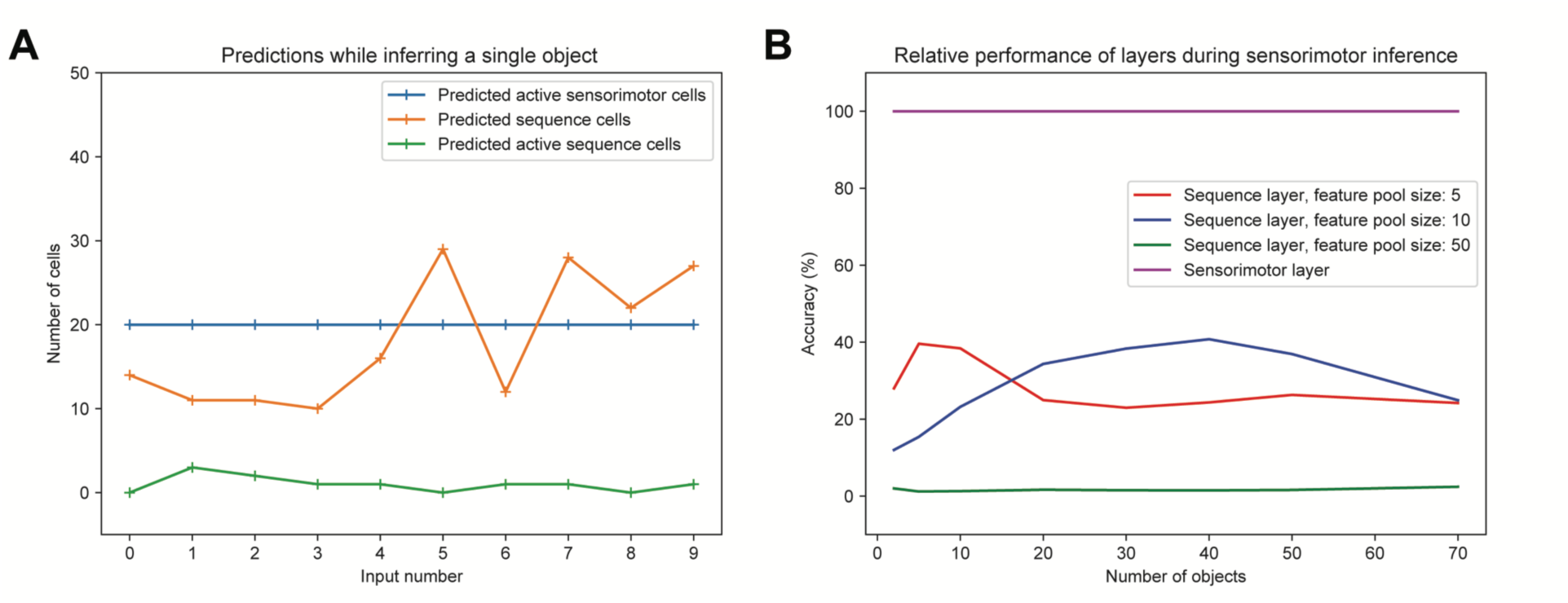
**A.** Example network behavior while inferring a single object. “Predicted” and “predicted active” have the same meaning as in the previous figure. A prediction of 20 cells which are all subsequently activated would constitute perfect prediction. The temporal sequence layer occasionally generates more than 20 predictions but fails to consistently make accurate predictions. As expected, the sensorimotor layer consistently makes perfect predictions. **B.** Relative accuracy of the sequence layer while exploring objects during sensorimotor inference. Each point represents the average prediction accuracy across all trained objects. For reference, we have included the performance of the sensorimotor layer, which achieves perfect accuracy in all conditions.

To better understand the behavior of the sequence layer half of the network during sensorimotor inference, we systematically varied the size of the feature pool and number of training objects. To quantify the performance, we calculated the fraction of input elements that were correctly predicted for each object. Since each object contains 10 features, chance performance is 10%. **Figure 5B** shows the results. With a small feature pool and a small number of objects, there is some chance of repeated feature transitions. The sequence layer has a higher likelihood of making predictions that are occasionally correct. With even a moderate feature pool size (e.g. pool size of 50 as in **Figure 5B**) this becomes extremely unlikely. Performance in this case is well below chance since the basal dendritic segments in the layer are simply not able to lock on to consistent repeated transitions in order to form any predictions. We verified that the sensorimotor layer achieved perfect accuracy in all these parameter combinations.

### Simulations with Combined Sequences

The previous simulations demonstrated the behavior of the model after training on either pure temporal or pure sensorimotor sequences. The experiments verified that the sensorimotor layer only makes reliable predictions for sensorimotor sequences in the presence of a statistically relevant location signal. Similarly, the temporal sequence layer only makes reliable predictions with pure temporal sequences. Next, we wanted to see how the network behaved when the stream of inputs contained a mixture both sequence types.

To do this we trained a single network on both types of sequences using the same pool of features. The training set consisted of 50 sequences (each of length 10) and 50 objects (each containing 10 points). The feature and location pools both had 50 elements. Sequences and objects were constructed as before. This is a challenging training set as there is significant sharing of features across objects and sequences as well as within them. In order to behave correctly, both layers must simultaneously form models that predict these shared features, but only if the predictions are statistically relevant under the appropriate context.

To test the network after learning, we presented a stream of inputs to the network, randomly intermixing sensorimotor and temporal sequences. As before, we measured the number of correctly predicted cells from each layer. **Figure 6** shows the behavior for a subset of the input stream. The layer with accurate predictions automatically switches depending on the underlying sequence type. Each layer occasionally makes extra predictions. This is understandable given the large number of shared feature elements. The figure shows that the network is able to disambiguate and separate temporal and sensorimotor sequences, even when both are composed of identical feedforward inputs.

**Figure 6.**
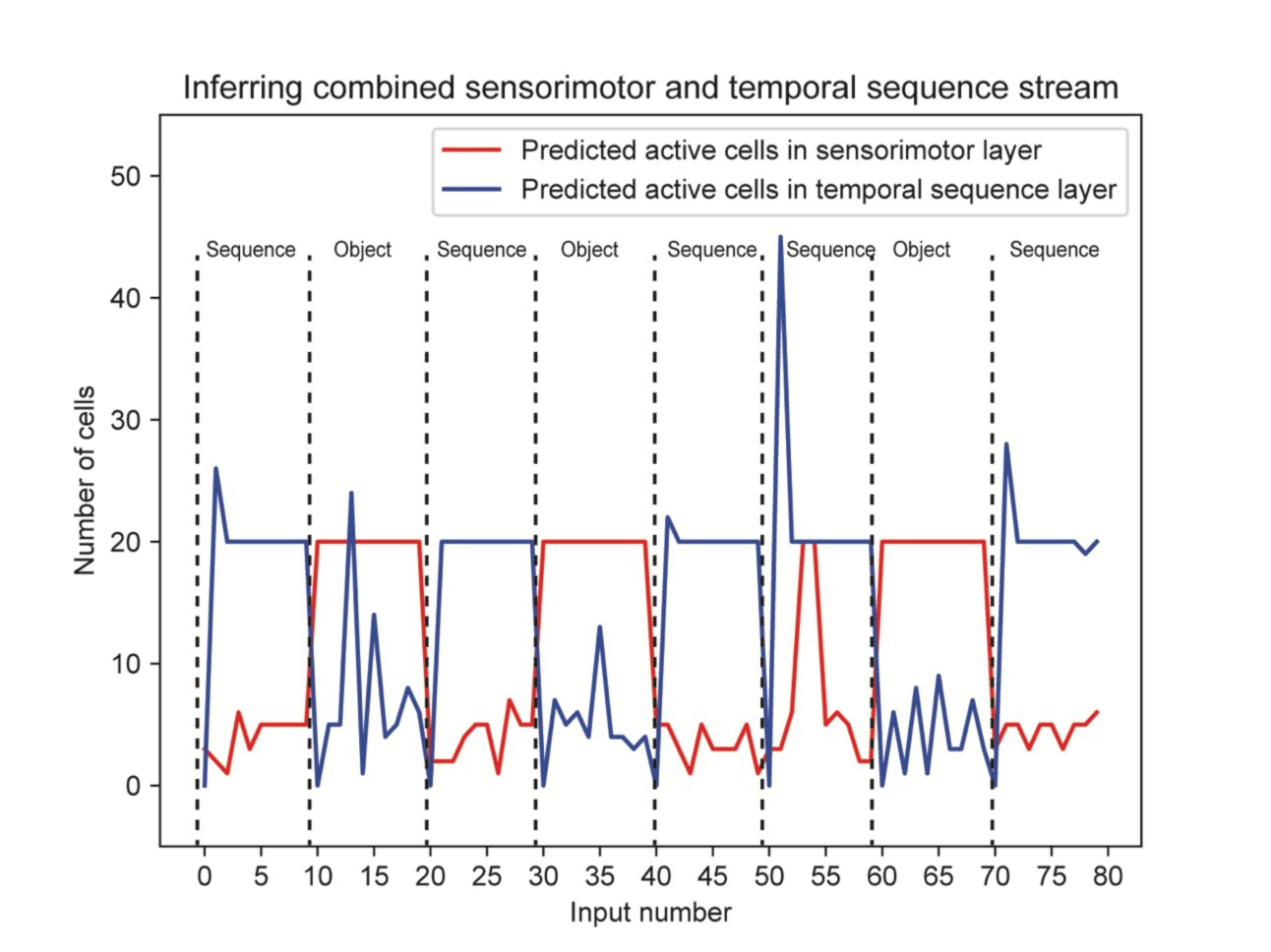
Network behavior while inferring a stream containing a mixture of temporal sequences (labeled “Sequence”) and sensorimotor sequences (labeled “Object”). The type of sequence was randomly changed every 10 inputs (dashed lines). The temporal sequence layer reliably recognizes temporal sequences (number of predictive active temporal cells equals 20). Similarly, the sensorimotor layer consistently recognizes objects. Both layers make occasional extra predictions. The temporal sequence layer occasionally contains substantially more than 20 predicted active cells. This is due to sequences with shared prefixes; a few timesteps are required to completely disambiguate a sequence.

## DISCUSSION

In this paper we reviewed a neural mechanism for prediction in cortex. The mechanism is based on a model of pyramidal neurons where contextual patterns are recognized on active dendritic segments which bias the cell towards firing. We described a network layer where detection of a contextual pattern leads to a sparse pattern of activity that is specific to that context. This arrangement is flexible. By using different sources of context, the network can learn different types of input transitions. When the context represents past activity within the layer, the layer forms a predictive temporal sequence memory. When the context represents locations derived from movement commands, the layer recognizes and predicts sensorimotor sequences. We then showed how a combined network, where the two layers share input and minicolumns, can learn both sensorimotor sequences and temporal sequences.

Sensory regions receive a sequence of inputs caused by a mixture of externally and internally generated events. Untangling sensorimotor sequences from externally generated sequences is an important problem that must be solved by the neocortex. We proposed a model that successfully achieved this task. Through simulations we explored the interactions between the diversity of input features, the diversity of behavioral motions, and the total number of objects and sequences learned by the network. Based on these results, we propose that the core of the solution lies in exploiting the underlying statistical predictability of movement based predictions vs. pure temporal sequences. We believe these represent a fundamental tradeoff in predictability that can be exploited by the cortex. With a sufficiently rich input stream and a sufficiently diverse range of motions it is possible to untangle a mixed input stream.

In (Hawkins and Ahmad, 2016) and (Hawkins et al., 2017) we list a number of predictions that are specific to the sequence memory and sensorimotor models respectively. These predictions include the emergence of highly sparse representations for predictable stimuli, denser representations for unpredictable stimuli, the specificity of cell assemblies to particular points in a sequence or object, predictions regarding branch specific plasticity, and predicted arrangements of fast spiking inhibitory cells. We have not discussed any new mechanisms in this paper, and as such, all those properties are valid here as well. The one new prediction from this paper is we expect to see all of the above properties, for both types of sequences, to be exhibited in a single population of cells (or cells in close proximity), see below.

### Mapping to cortical anatomy

Where might the network described in this paper exist in the neocortex? If both types of sequence processing (sensorimotor and pure temporal) occur in all sensory modalities, then we would expect to see the neural circuitry that supports them to be present in all sensory regions.

In (Hawkins et al., 2017) we proposed that the primary input layer, L4 was the sensorimotor inference layer. The following properties suggest this assignment. Cells in L4 are driven by feedforward input to the region, L4 cells in vertical alignment have similar feedforward receptive fields (Favorov and Whitsel, 1988; Mountcastle, 1997; Jones, 2000; Buxhoeveden, 2002). The receptive fields of upper layers diverge in different contexts (Allman et al., 1985; Vinje and Gallant, 2000). Interestingly, approximately half of the synapses on L4 basal distal dendrites come from L6a (Ahmed et al., 1994; Binzegger et al., 2004), which is a logical candidate for conveying the location input. We also argued that the object layer in our model is located in L2 and/or L3. This is suggested because L4 cells are drivers to L2/3 cells (Lohmann and Rörig, 1994; Feldmeyer et al., 2002; Sarid et al., 2007). L2/3 cell axons leave the region and thus constitute one of the primary outputs of a region (Douglas and Martin, 2004), and L2/3 cell receptive fields are more stable than L4 cell receptive fields (Gur and Snodderly, 2008). Another reason is that L2/3 cells have long range intra-laminar connections which support inter-column voting and faster inference (Bosking et al., 1997; Stettler et al., 2002; Hunt et al., 2011). For these reasons, we continue to propose L4 is the sensorimotor input layer and L2/3 is the object layer.

Where then would the temporal sequence layer reside? We suggest two possibilities. We don’t have sufficient data to eliminate either. The first possibility is that L4 acts as both a sensorimotor and a temporal sequence layer. L4 cells connect to each other (the requirement for temporal memory) (Feldmeyer et al., 1999) and they receive a large input from L6a (the requirement for sensorimotor memory). Although it is convenient to think about sensorimotor and pure sequence memory as separate processing layers we don’t see why this has to be the case. In our simulations we segregated the neurons within a minicolumn by context type, but we see no reason that the network would perform differently if the contextual inputs were mixed together.

The second possibility is that L3b is the temporal sequence layer. Like L4, lower L3 cells receive direct sensory input (White and Hersch, 1981; Thomson and Lamy, 2007) which is consistent with L3b being the temporal sequence layer. Unlike L4, L3b cells are not known to receive a strong input from L6a. The largest input to distal basal dendrites in L3 cells is from other cells in L3, which is consistent with the context needs of temporal sequence memory. If it was found that L3b cells get direct feedforward input AND send their axons outside of the region (Markov et al., 2014), then this would be inconsistent with L3b being temporal sequence memory. Further anatomical data would help resolve this uncertainty.

### Physiological mechanisms

There is now a wealth of experimental evidence that primary sensory areas can learn and model sequences (Vinje and Gallant, 2000; Gavornik and Bear, 2014; Miller et al., 2014; Pitas et al., 2016). The exact neural mechanisms for sequence learning in cortex are still unknown. Our model proposes that active dendrites using simple learning rules is a core mechanism for learning complex temporal transitions. In (Hawkins and Ahmad, 2016) we reviewed in detail the biophysics of active dendrites as it relates to this hypothesis.

(Chait et al., 2007) provides evidence that auditory cortex can optimally model orderly sequence transitions, and differentiate from unpredictable transitions. They propose the existence of "a general mechanism that operates early in the processing stream on the abstract statistics of the auditory input". (Cavanaugh et al., 2016) provides evidence that efference copy is required for visual perceptual stability in the context of eye movements. In a way, the mechanism presented here is a generalization of these two core ideas. Our model shows how efference copy can provide the statistical structure required for separating sensorimotor sequences from pure temporal sequences.

In motivating our model, we have discussed the importance of context in forming the right predictions. (Phillips, 2015) discusses the biophysics of incorporating context. The paper describes several intracellular and active dendritic mechanisms for amplifying and modulating a single neuron's response, and their impact on cognitive phenomena. Although the paper does not specifically consider temporal sequences, the mechanisms described (such as the impact of apical dendrites) form a possible biophysical basis for contextual modulation in our model.

### Models of sequence memory

Sequence learning has been studied extensively in the field of machine learning. These algorithms don’t attempt to model the biology but focus on generic sequence learning tasks. It might in theory be possible to extend those algorithms to model combined sensorimotor sequences. However existing machine learning methods for robot sensorimotor learning (Pinto et al., 2016; Devin et al., 2017) are significantly different from machine learning techniques for temporal sequences (Waibel, 1989; Hochreiter and Schmidhuber, 1997; Fine et al., 1998). A detailed comparison of sequence learning techniques to our model can be found in (Cui et al., 2016; Hawkins and Ahmad, 2016).

A comprehensive review of biologically motivated sequence memory models can be found in (Dehaene et al., 2015). The paper describes a rich taxonomy of different sequence types, representing increasing levels of complexity. Two points are worth emphasizing here. First, they do not include sensorimotor sequences in their taxonomy. We have suggested that these sequences contain a statistical structure that is different from that of pure temporal sequences. As such, sensorimotor sequences might represent a different category of sequences altogether.

Second, they discuss the importance of modeling specific time intervals, such as the timing of notes in a song (Memmesheimer et al., 2014). Sensitivity to the exact timing of inputs is a general capability that is applicable to both sensorimotor sequences (Crowe et al., 2014) and pure temporal sequences (Gavornik and Bear, 2014). We have not described how our model could represent the precise timing of sequence elements, but a natural extension would be to include timing signals as additional contextual input. Such signals are available to the cortex through various pathways (Crowe et al., 2014; Mello et al., 2015). Incorporating these signals could allow networks of HTM neurons to model specific time delays, and represents an interesting area for future work.

(Kastellakis et al., 2016) discuss a recent model of associative memory that involves synaptic clustering on active dendritic branches. They show that memories can be linked across time. Their model is related to the one presented here in that they use active dendrites where dendritic segments can represent transitions in a sequence. The model focuses on understanding some of the biophysical properties of synaptic plasticity and memory capture. Currently the model is not designed to represent complex sequences of the type presented here, and it has not been applied to sensorimotor inference. It might be possible to implement extensions that use minicolumns or the inhibitory mechanisms in our model.

## MATERIALS AND METHODS

Details of the methods presented here can be found in (Cui et al., 2016; Hawkins and Ahmad, 2016; Hawkins et al., 2017). Here we list the data and simulation details that are specific to this paper.

### Data generation details

For the pure temporal sequence experiments we generated sequences of length 10. We generated each temporal sequence by creating a set of 10 features chosen randomly with replacement from a fixed pool of features. Thus the same feature could appear multiple times within a sequence, and in different sequences. Accurate prediction therefore required the system to maintain past temporal context.

For sensorimotor sequences we generated a set of objects as in (Hawkins et al., 2017). Each object was comprised of 10 points. Each point consisted of a feature and location vector randomly chosen from two corresponding pools of vectors. Features were chosen randomly with replacement whereas locations were chosen without replacement. Any given object could have multiple points with the same feature.

Although there could be significant sharing of features across objects and sequences, we ensured that each object and sequence was unique for any given trial. The network was reset at the end of each sequence or object presentation. The reset zeroed out activity in the layers and placed the network into a canonical state before the next sequence/object was presented.

Each feature vector consisted of a binary vector with 512 components (one per minicolumn), of which a random 20 positions were set to 1. Each location vector was a binary vector with 1024 components, of which a random 20 positions were set to 1.

### Simulation details

As with our previous papers, we relied on the NuPIC codebase to implement the simulations in Python and C++. For this paper we used the same code implementation to model the sensorimotor and temporal sequence layers, specifically the *ExtendedTemporalMemory* class. We used identical layer sizes and learning parameters for both layers as follows:

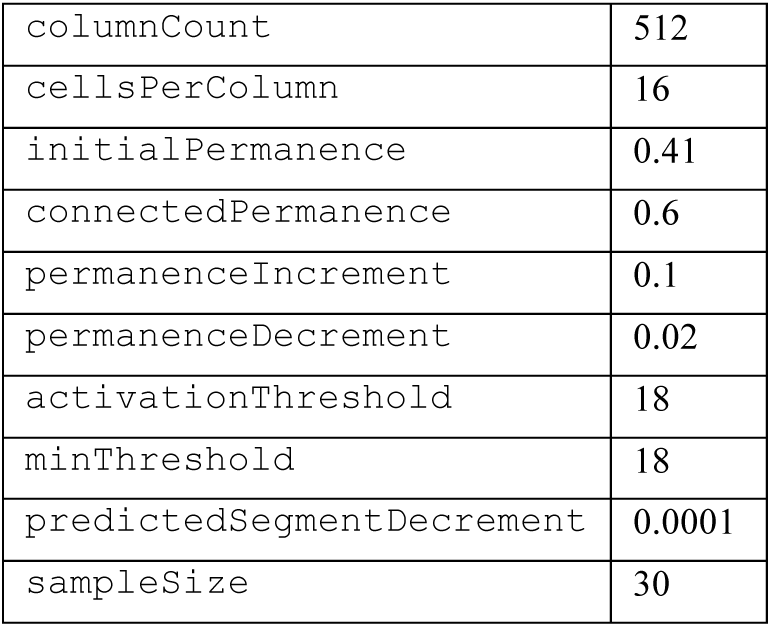

|  |  |
| --- | --- |
| columnCount | 512 |
| cellsPerColumn | 16 |
| initialPermanence | 0.41 |
| connectedPermanence | 0.6 |
| permanenceIncrement | 0.1 |
| permanenceDecrement | 0.02 |
| activationThreshold | 18 |
| minThreshold | 18 |
| predictedSegmentDecrement | 0.0001 |
| sampleSize | 30 |

The primary difference was the source for the context vectors, as described in the text. The source files for the simulations are available on Github at: https://github.com/numenta/htmresearch/projects/combined_sequences/.

## ACKNOWLEDGEMENTS

We thank Scott Purdy, Yuwei Cui, Marcus Lewis, and numerous other collaborators at Numenta for many discussions that have improved the ideas in this paper.

